# The role of RNA in the maintenance of chromatin domains as revealed by antibody mediated proximity labelling coupled to mass spectrometry

**DOI:** 10.1101/2023.10.16.562458

**Authors:** Rupam Choudhury, Anuroop Venkateswaran Venkatasubramani, Jie Hua, Marco Borsò, Ignasi Forne, Axel Imhof

**Affiliations:** Department of Molecular Biology, Biomedical Center Munich, Ludwig-Maximilians University, Großhaderner Strasse 9, 82152 Planegg-Martinsried, Germany; Graduate School of Quantitative Biosciences (QBM), Ludwig-Maximilians-Universität München, Feodor-Lynen-Strasse 25, 81377, Munich, Germany; Protein Analysis Unit, Biomedical Center (BMC), Faculty of Medicine, Ludwig-Maximilians-University (LMU) Munich, Großhaderner Straße 9, 82152, Martinsried, Germany

## Abstract

Eukaryotic chromatin is organised in individual domains, which are defined by their location within the nuclear space, the molecular interactions within each other and their distinct proteomic compositions. In contrast to cellular organelles surrounded by lipid membranes, the composition of distinct chromatin domains is rather ill described and highly dynamic. To identify proteins associated with a specific chromatin domain and to better understand how those domains are established and maintained, we used a new method, which we termed AMPL-MS (Antibody mediated proximity labelling mass spectrometry). This method is based on antibodies and does not require the expression of fusion proteins and therefore constitutes a versatile and very sensitive method to characterize chromatin domains containing specific signature proteins or histone modifications. We used AMPL-MS to characterize the composition of chromocenter as well as the chromosome territory containing the hyperactive X-chromosome in *Drosophila*. As an outcome of this method, we show the importance of RNA in maintaining the integrity of these domains as treatment with RNAse alters their proteomic composition. Our data demonstrate the power of AMPL-MS to characterize the composition of non-membranous nuclear domains.

## Introduction

Genetic information in the eukaryotic nucleus is packaged in a complex chromatin structure. Typically, chromatin has been classified in two major categories: euchromatin, which is loosely packed and more accessible, and heterochromatin, which is tightly packed and less accessible ^1,2^. These different degrees of accessibility are thought to be mediated by a network of specific protein-DNA and protein-protein interactions^3^. Systematic mapping studies of proteins on chromatin using DamID^3,4^ or ChIP^5^ revealed a diverse composition of chromatin resulting in up to 30 different types of chromatin. A variety of different chromatin capture methods have demonstrated that the genome can be separated in two large functional compartments^6^ with inter-chromosomal interactions occurring mainly between loci belonging to the same compartment. While it is generally accepted that the three-dimensional organization of nuclear chromatin in distinct domains plays a major role in regulating gene expression^7–9^, the mechanisms of how the different types of chromatin form and how they contribute to the dynamic regulation of gene expression are far from being understood. Several studies have shown that specific chromosomes or classes of chromatin occupy distinct areas within the nucleus, often called territories^10,11^ or domains^12^. Among the most prominent examples of such regions are the centromeric heterochromatin that frequently localizes at the nuclear periphery^13^ or cytologically identifiable nuclear bodies like the nucleolus that is formed on the rDNA locus^14^ or Cajal bodies that are linked with snRNP and snoRNP biogenesis^15^. These nuclear bodies and the distinct chromosome territories dissolve and re-form in G1 upon conclusion of mitosis^16,17^. Such self-organized formation and maintenance of functional domains is most likely mediated by multiple interactions between the DNA, proteins and RNA found within these domains^18^. The individual interactions that drive this process are often weak in nature but nevertheless able to drive the formation of distinct nuclear bodies. Though microscopically detectable, many nuclear bodies are highly dynamic and difficult to purify. In fact, even for the most abundant classes of nuclear bodies such as nucleoli or cajal bodies, the investigation of their proteomic composition required large quantities of cultured cells^19–21^. The purification and characterization of specific chromosomal territories is even more challenging and often involves the disruption of the nuclear 3D structure before purification of a chromosomal domain^22–24^. Therefore, these purification methods frequently depend on the stable interaction of the protein with the DNA and hence many weak interactions are lost. A possible solution to this loss is the use of proximity biotinylation methods ^25^, which have been shown to provide powerful tools to identify and characterize such weak interactions^26,27^. In fact, proximity biotinylation has been increasingly used in chromatin research ^28–32^ to characterize the chromosomal environment of DNA bound factors that have been elusive to ChIP or ChIP-MS methods. The disadvantage of most methods used so far is the fact that proteins needed to be expressed as fusion proteins with BioID or variants thereof or with Apex2 and new cell lines had to be established^28,29,33,34^. Moreover, as the expression of transgenic fusion proteins is difficult to control and the delivery of biotinylation reagents is hard to control, it results in a high background and variance of the proximity proteome. To overcome these issues, various in vitro methods using BirA derivatives^30,31,35^ have been recently established to facilitate proximity biotinylation in cells. The disadvantages of BirA and its derivatives are their relatively low biotinylation efficiency and slower kinetics resulting in a lower sensitivity and the requirement of large amounts of input material. We therefore developed and tested a pipeline using a proteinA-Apex2 fusion protein and tested its efficiency on specific chromosomal domains in *Drosophila*. This method, which we call AMPL-MS for Antibody mediated proximity labelling coupled to mass spectrometry, has so far only been used to study a limited degree of histone modifications^36^. The extension of this method towards various nuclear proteins and its usage to study the changes upon a challenge enabled us to investigate specific chromatin domains in insect tissue culture cells and demonstrate the importance of RNA in the formation of these domains.

## Results

### Versatile proximity biotinylation in cells

To establish a versatile and sensitive method to characterise the proteomic neighbourhood of a given chromatin protein, we expressed a His-tagged fusion protein of Protein A and Apex2 in *E. coli* (pA-Apex2; Supplementary Fig. S1). The fusion protein was used to tether Apex2 to chromatin domains marked by specific antibodies, which allows an efficient biotinylation of all proteins in the vicinity (Fig. 1a). To assess the specificity of the method, we first targeted centromeric chromatin using an antibody against the centromeric histone variant of H3, CID (dCenpA). The colocalization of the biotin signal with CID in immuno-fluorescence images shows that pA-Apex2 biotinylates proteins within the centromeric domain in presence of an anti-CID antibody, biotin phenol and hydrogen peroxide (Fig. 1b,c). Proteins within this domain were then isolated using streptavidin beads and analysed by mass spectrometry leading to the identification of 172 proteins that localized in proximity to CID containing centromeric chromatin (Fig. 1d). All previously characterized centromeric proteins of *Drosophila* were almost exclusively detected in proximity to CID (Fig. 1e)^28,37^. A gene ontology (GO) enrichment analysis of the proteins that had not yet been reported as CID interactors revealed a strong enrichment of factors involved in RNA related processes (Fig. 1f). Due to the much lower background and the increased sensitivity of the AMPL-MS method we got a comparable number of identified components of centromeric chromatin domain from as little as 2×10^7^ cells as opposed to 2 × 10^10^ cells when expressing CID-Apex2 ^28^ (Supplementary Fig. S2d,e). The establishment of AMPL-MS as a sensitive and versatile method to analyse chromatin domains allows a quick investigation and comparison of such domains even in difficult to isolate cell populations.

**Fig. 1.**
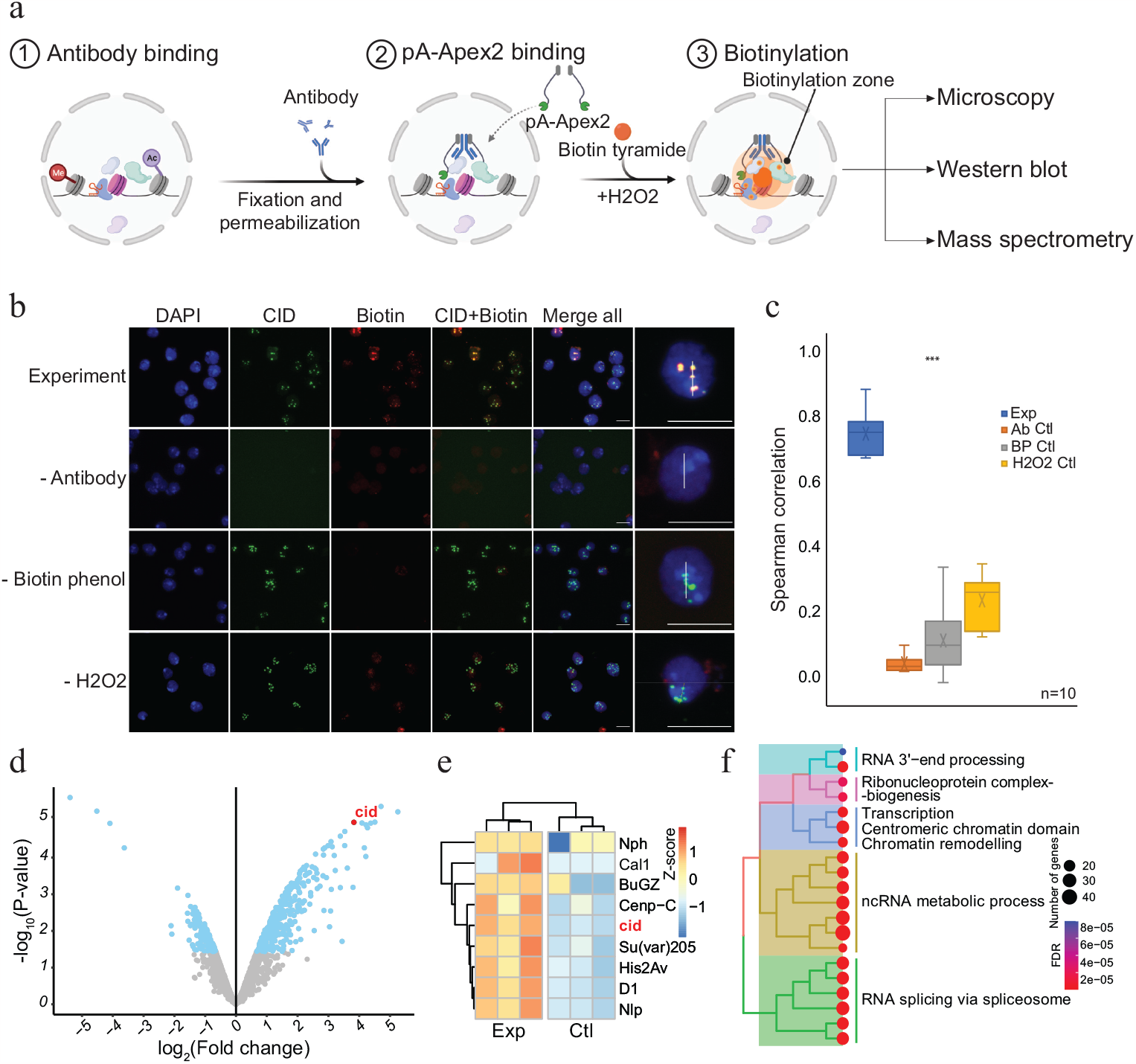
AMPL-MS to study the proteomic composition of the Drosophila centromere in a genetically unperturbed cell line. **(a)** Schematic of AMPL-MS. Isolated nuclei were fixed, permeabilized and incubated with specific antibodies. Recombinant pA-Apex2 enzyme binds to the antibody and biotinylates associated proximal proteins upon the addition of H_2_O_2_ and biotin phenol. Proximal protein biotinylation is visualised by microscopy, western blot and proteins were identified by mass spectrometry. The schematic **(b)** Immunofluorescence microscopy of centromeres using a Cenp-A (CID) antibody (in green) and the corresponding proximity proteome after biotinylation by pA-Apex2 (in red). Nuclear DNA was stained by DAPI (in blue). IgG was used as antibody control. Scale bars represent 10 μm (large panel) or 5 μm (small panel). **(c)** Distribution of pair-wise Spearman correlations for quantifying the relationship between CID and biotinylation. Images of 10 cells from three independent experiments were used. Statistical significance is based on Wilcoxon rank sum test (***P-value< 0.001). **(d)** Volcano plot of purified biotinylated proteins identified by mass spectrometry. The bait protein is highlighted in red. The x-axis represents the log2 fold change and the y-axis represents -log10p-value comparing three IgG replicates with three CID antibody replicates (paired). The significantly enriched proteins (abs(LFC)>1 and padj≤0.01) are highlighted in blue. **(e)** Heatmap showing the enrichment of known centromeric proteins. The heatmap was plotted using scaled log2 raw intensities. Each column represents values obtained from three independent biological replicates. **(f)** Over representation analysis showing top 20 biological processes (BP) using significant proteins from **(d)**. Unsupervised clustering was performed for the GO terms. The colour gradation form blue to red represents FDR (False Discovery Rate) and dot size represents the number of proteins found enriched in the named pathway (count).

### AMPL-MS allows the distinction of different chromatin domains

To assess the applicability of AMPL-MS to differentiate between different chromosomal territories, we wanted to characterize the proteomic neighbourhood of the transcriptionally hyperactive X-chromosome in *Drosophila*. To do this we used an antibody recognising MSL2, a component of the dosage compensation complex that specifically and selectively associates with the male X chromosome^38^. Like the selective biotinylation of the centromere, AMPL-MS using an anti-MSL2 antibody resulted in the specific labelling of the X chromosome territory (Fig. 2a,b). As expected, the composition of the enriched sets of protein depends on the bait identity (Fig. 2c and Supplementary Fig. S3). Known centromeric proteins were more enriched in the anti-CID AMPL-MS proteome and proteins known to localize the hyperactive X chromosome were mainly detected in the anti-MSL2-AMPL-MS proteome (Fig. 2d). The analysis of the entire dataset by a principal component analysis also showed a tight clustering of the domain proteome according to the bait identity (Supplemental Fig. S2d). A gene ontology (GO) enrichment analysis of the differentially enriched proteins revealed a bias for the GO terms spindle assembly and chromatid segregation in proteins close to CID and sex determination and dosage compensation in the proteins proximal to MSL2. Interestingly, both chromatin neighbourhood contained factors related to RNA related processes albeit with slightly different functional terms (Fig. 2e,f). As both proteins are tightly associated with chromatin, we wondered whether the histones in proximity to CID carry different modification patterns than the one in the neighbourhood of msl2. In agreement with the function of the bait proteins we detect a moderate increase of activating marks in the MSL2 proximity proteome and a slightly higher level of repressive marks in proximity of CID (Fig. 2g). These comparative experiments show that we can apply AMPL-MS to distinguish distinct chromatin domains based on their proteomic composition.

**Fig. 2.**
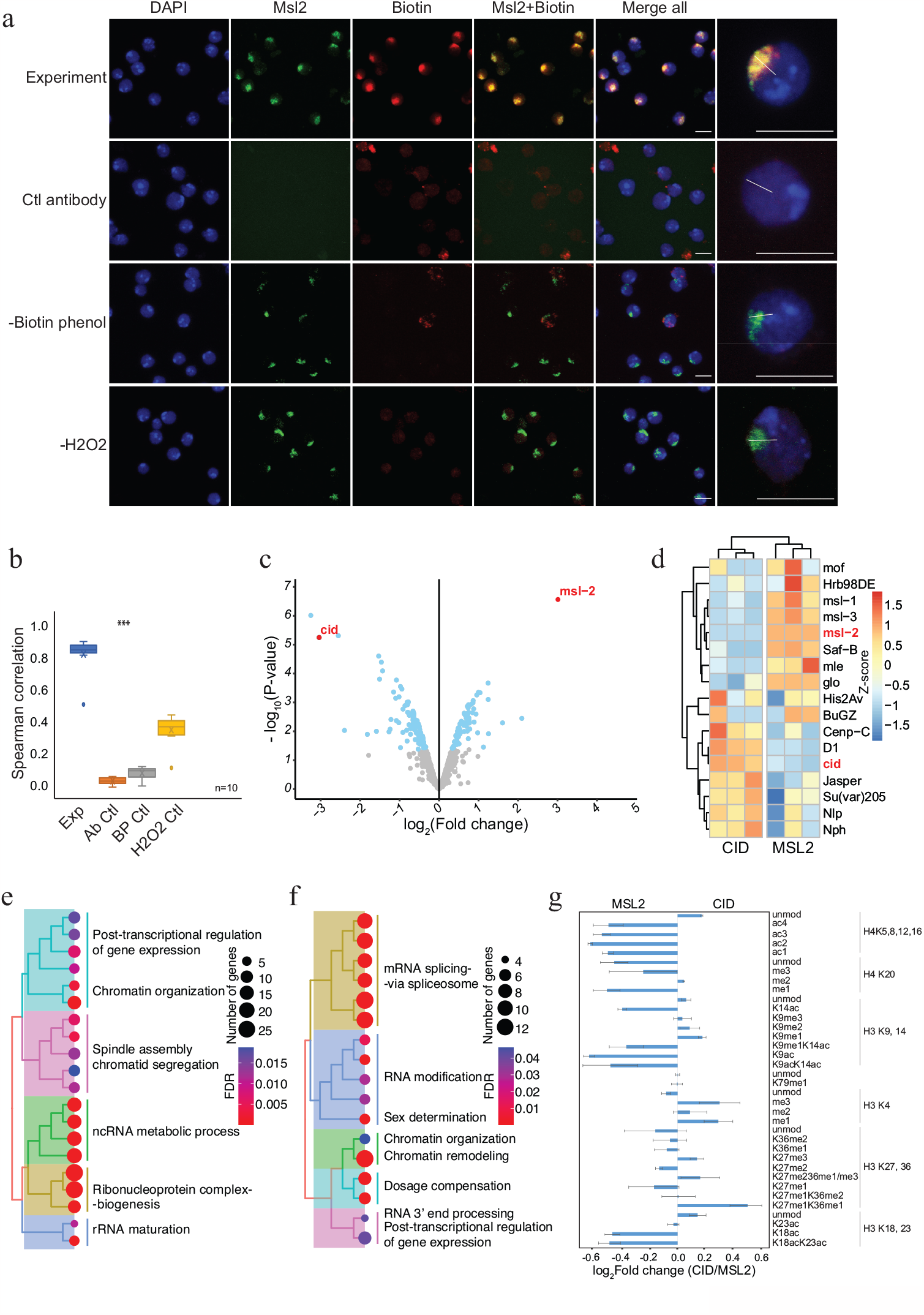
AMPL-MS can efficiently identify protein associated with different chromatin domains. **(a)** Immunofluorescence of the X-chromosome bound by MSL2 antibody (in green) and biotinylated proteins after biotinylation by pA-Apex2 (in red). Nuclei were stained by DAPI (in blue) and IgG was used as antibody control. Scale bars represent 10μm (large panel) or 5μm (small panel). **(b)** 10 cells from three independent experiments were used to quantifying the relationship between MSL-2 and biotinylation. Images of 10 cells from three independent experiments were used. Wilcoxon rank sum test is used for comparison (***P-value< 0.001). **(c)** Volcano plot of biotinylated proteins identified by mass spectrometry. The bait proteins are highlighted in red. The x-axis represents the log2 fold change and the y-axis represents -log10p-value comparing three CID replicates with three Msl-2 antibody replicates (paired). The significantly enriched proteins (LFC>1 and padj≤0.01) are highlighted in blue. **(d)** Heatmap displaying the enrichment of known centromeric and X-chromosome associated proteins. The heatmap was plotted using scaled log2 raw intensities. Each column represents values obtained from three independent biological replicates. Over representation analysis showing top 20 Biological process (BP) for significantly enriched proteins from **(c)** associated with either centromere **(e)** or the X-chromosome **(f)**. The colour gradation form blue to red represents FDR (False Discovery Rate) and dot size represents the number of proteins found enriched in the named pathway (count). **(g)** Relative quantification of histone modifications associated with the X-chromosome territory (left) and the centromere (right).

### Proximity labelling of proteins associated with post-translationally modified histones

Having established AMPL-MS as an efficient and sensitive method to identify proteins associated with distinct chromatin domains marked by antibodies that recognize signature proteins, we wanted to test whether we could also use AMPL-MS to characterize the proteome in proximity to specific histone marks. To this end, we used antibodies recognising H3K4me3, H3K9me3 and H4K16Ac for AMPL-MS. Consistent with the role of these histone modifications in establishing transcriptional active (H3K4me3, H4K16ac) or repressive (H3K9me3) chromatin domains, the proteomic composition of these domains are very different (Fig. 3a and Supplementary Fig. S4). As expected, proteins in proximity to H3K9me3 includes several known K9me3 binding proteins, such as Su(var)205 (HP1a) or HP5 and other known heterochromatin associated proteins such as Su(var)3-7, Su(var)3-3, HP5, HDAC3, HDAC1, HDAC6 (Fig. 3b). Interestingly, we also find an overlap with the centromeric H3 variant Cid and proteins that we detected in proximity of it like NPH, NLP or CENP-C. In contrast, we find mainly proteins associated with active chromatin and the components of the dosage compensation complex in proximity of H4K16ac (Fig. 3b).

**Fig. 3.**
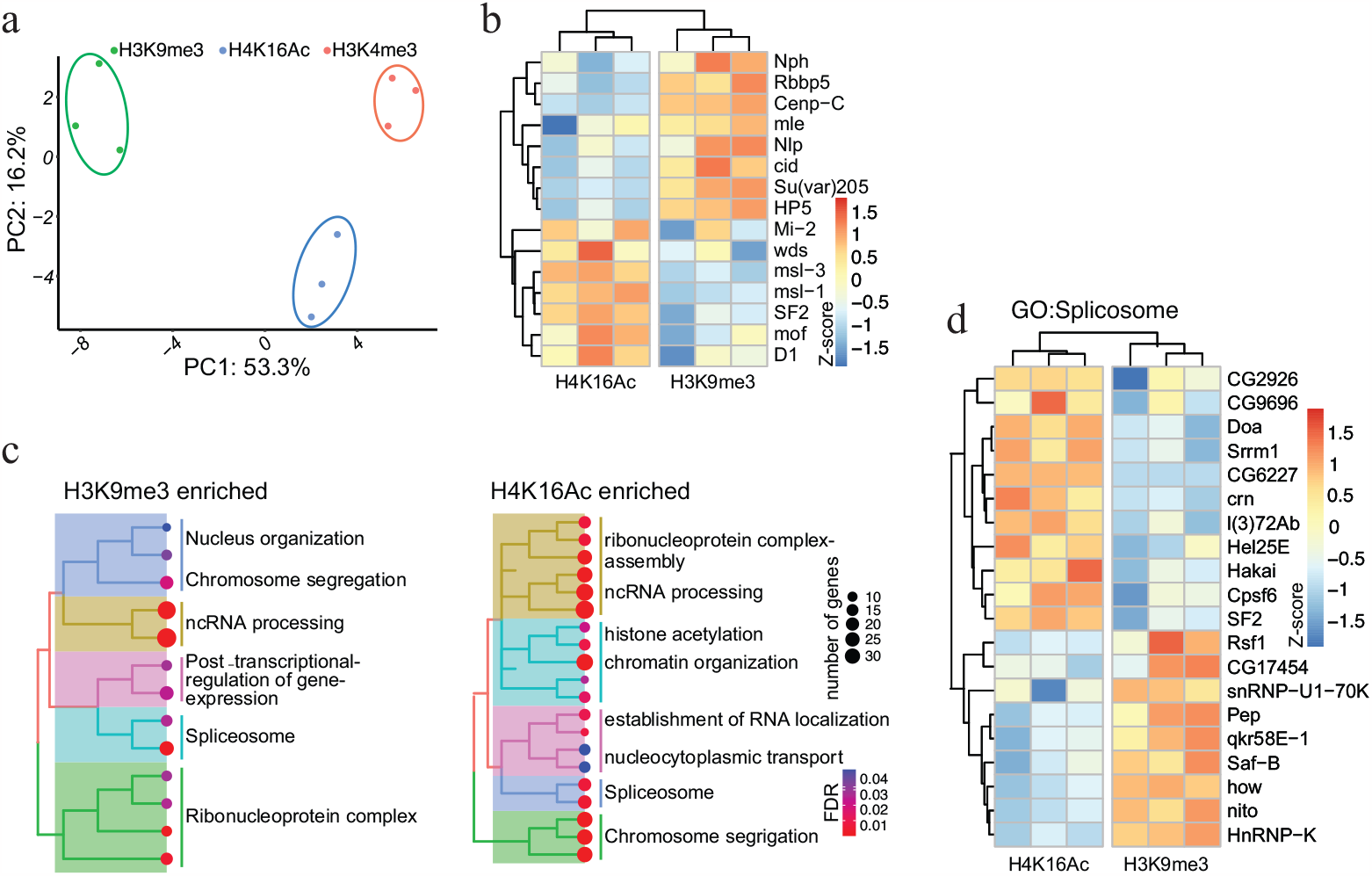
Protein associated with different and dynamic histone modifications can be effectively identified by AMPL-MS. **(a)** Principal Component Analysis (PCA) based on the proteomic composition for three different histone marks: H3K9me3, H4K16ac, and H3K4me3 antibodies. **(b)** Heatmap displaying the enrichment of known H3K9me3 and H4K16ac histone modification associated proteins. The heatmap was plotted using scaled log2 raw intensities. Each column represents values obtained from three independent biological replicates. **(c)** Over representation analysis showing top 20 Biological process (BP) for the significantly enriched protein associated with either H3K9me3 or H4K16ac histone modifications. The colour gradation form blue to red represents FDR (False Discovery Rate) and dot size represents the number of proteins found enriched in the named pathway (count). **(d)** Heatmap for protein categorized under the GO term “Spliceosome”, a common GO term found for proteins associated with H3K9me3 and H4K16ac histone modification. The heatmap was plotted using scaled log2 raw intensities. Each column represents values obtained from three independent biological replicates.

Similar to the comparative analysis of CID and MSL2 we find a number of factors involved in splicing and RNA processing in the neighbourhood of H3K9me3 or H4K16ac containing chromatin (Fig. 3c). Despite having the same gene ontology term associated with, the individual proteins in proximity to H3K9me3 or H4K16ac were quite different (Fig. 3d). Factors like SAF-B or HNRNP-K, which we detect in proximity to H3K9me3 have been shown to play an RNA dependent role in heterochromatin organisation^39^, in Xist mediated transcriptional silencing^40^. Several RNA binding proteins we detect closer to H4K16 on the other hand are part of the canonical or non-canonical splicing machinery^41^ and are often found close to sites of active transcription. The abundance of specific RNA binding proteins in proximity to various chromatin domains suggests a major and specific role of RNA in the organisation of chromosomal domains.

### The removal of RNA changes the proteomic environment of chromatin domains

Chromatin associated RNA has been suggested to serve as an architectural component or by facilitating the formation of membraneless condensates within the nucleus^42–47^. Many of the RNAs can act in cis as well as in trans and often show a rather specific distribution. For instance, dosage compensatory RNA Xist in mammals or Rox in *Drosophila* specifically associate with the inactive female X or the hyperactive male X-chromosome in vivo^48–51^. RNA transcribed from pericentromeric or centromeric chromatin bind plays a key role in setting up the structure of the centromere and the clustering of distinct centromeres in interphase^52,53^. To investigate the RNA dependent proteomic neighbourhood of distinct nuclear domains like the hyperactive X-chromosome or the chromocenter, we performed AMPL-MS for MSL2 and CID in the presence or absence of RNase A (Fig. 4a). As previously shown, RNA depletion disrupts the centromeric domain and the hyperactive X-chromosome, while maintaining the overall nuclear morphology (Fig. 4b,c). Thanks to the AMPL-MS method we could study the effect of an RNaseA treatment on the proteomic environment of the signature proteins of chromosomal domains. As expected, neither the targeted signature factor or proteins that mainly interact with them protein-protein interactions such as MSL1,3 and MOF for MSL2 or Cenp-C for Cid are not affected by RNAase treatment. However, the proteomic composition of the characterised domains changes substantially upon RNAse treatment (Fig. 4d,f). Not surprisingly, a large percentage of proteins that depend on the presence of RNA contain RNA binding domains (Fig. 4e,g), suggesting that the proximity is mediated by a direct interaction of these factors with RNA. While we have not identified the RNAs responsible for the formation of theses domains, we clearly observe that they do confer specificity for the domains as we observe very little overlap in the factors lost from the corresponding domains (Fig 4h). Consistent with their respective chromatin state we find repressive factors such as SAF-B or HNRNP-K losing their proximity to centromeric chromatin and activating factors such as the TAFs or components of the brahma complex moving away from the hyperactive X-chromosome. Interestingly, among the few common proteins that are lost from both domains is the RNA helicase MLE, which has been shown to influence dosage compensation^50,54,55^ as well as several other nuclear processes such as splicing and heterochromatin deposition^56^. The vertebrate homologue of MLE, DHX9, has been shown play a key role in the formation of RNA-DNA hybrids (R-loops) ^57^ which have been suggested to play an important role in centromere function in other organisms ^58–60^. Based on the observation that MLE can be detected within the chromocenter of Drosophila polytene chromosomes ^61^, a potential role for MLE for centromere formation in Drosophila has also been proposed. In fact, we also observe a minor fraction of MLE close to the centromere, which is lost upon RNase treatment (Supplemental Fig. S5), supporting this hypothesis (Fig 4i).

**Fig. 4.**
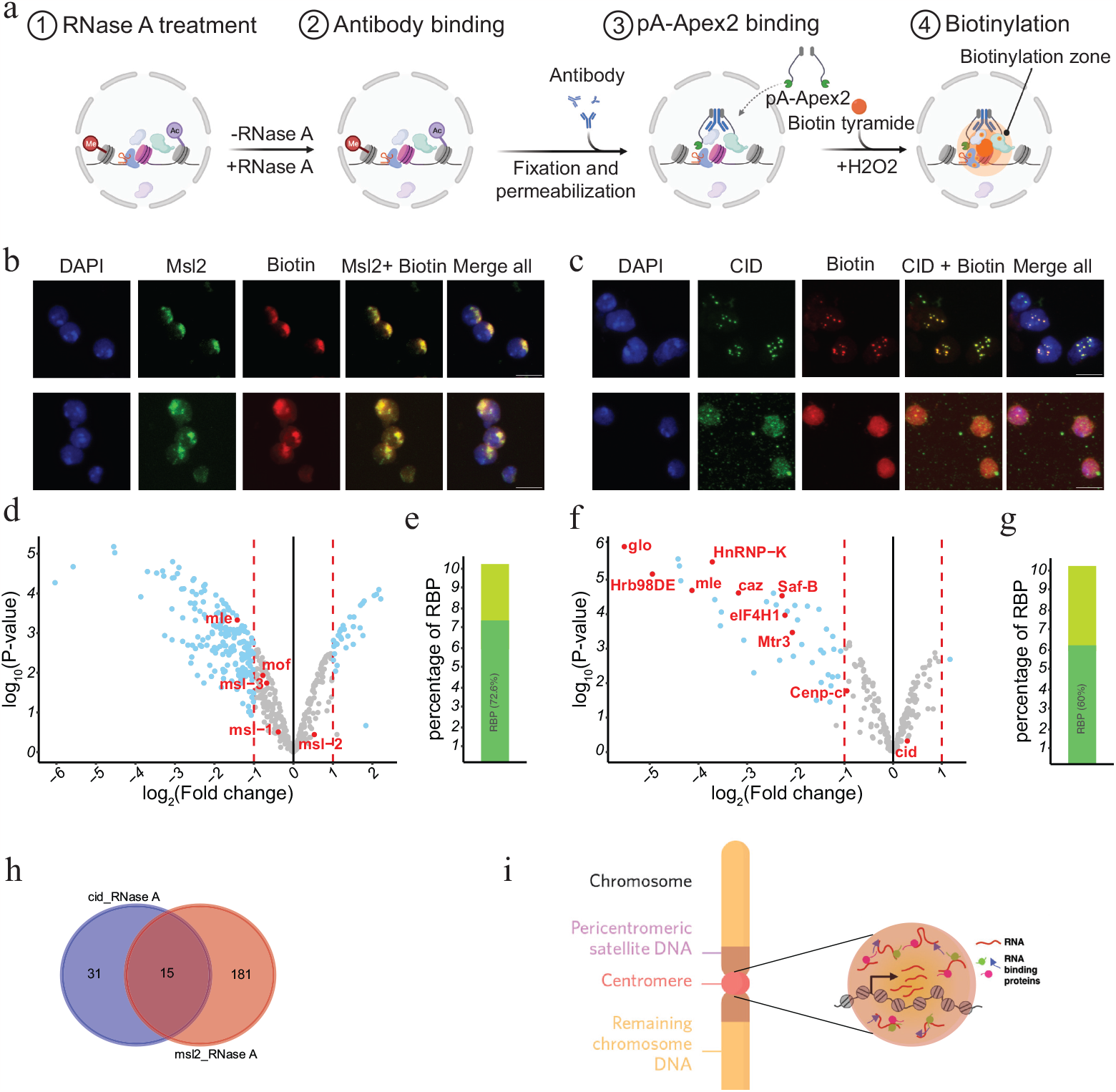
RNase treatment changes the proteomic environment of nuclear domains. **(a)** Schematic of AMPL-MS-RNase method. Isolated nuclei were treated with RNase A, fixed, permeabilized and incubated with specific antibodies. Recombinant pA-Apex2 enzyme binds to the antibody and biotinylates associated proximal proteins upon the addition of H_2_O_2_ and biotin phenol. **(b)** Immunofluorescence of the X-chromosome bound by a Msl2 antibody (in green) and biotinylated proteins after biotinylation by pA-Apex2 in control (i) and RNase A treated samples (ii). Nuclei was stained by DAPI (in blue). Scale bars represent 5μm. **(c)** Immunofluorescence of the centromer bound by an anti-Cid antibody (in green) and biotinylated proteins after biotinylation by pA-Apex2 in control (i) and RNase A treated samples (ii). Nuclei were stained by DAPI (in blue). Scale bars represent 5μm. Volcano plot of proteins identified by AMPL-MS-RNase using an anti Msl2 (**d**) or an anti-Cid antibody (**e**). The bait protein is highlighted in red, along with components of the drosophila dosage compensation complex. The x-axis represents the log2 fold change and the y-axis represents -log10p-value comparing three control MSL-2 AMPL-MS replicates with three MSL2-RNase A treated AMPL-MS replicates (paired). The significantly enriched proteins (LFC>1 and padj≤0.01) are highlighted in blue. **(f)** Overlap of proteins associated with X-chromosome and the centromere domain in an RNA dependent manner. **(g)** A schematic representation of RBPs associated with the centromere in an RNA dependent manner. The RNA molecules present at the centromere can recruit chromatin factors to form centromeric domain.

In summary our data show that the versatility of AMPL-MS allows us to characterize membrane-less domains in the nucleus at an unprecedented molecular level and therefore identify systematic changes in these organelles upon challenge.

## Methods

### Cell culture

*Drosophila* S2DGRC or L2–4 cells were grown in Schneider medium supplemented with 10% fetal calf serum, penicillin and streptomycin at 26°C.

### Cloning

To construct a bacterial expression vector for protein-A-Apex2-6xHis fusion protein (pk19pA-Apex2), the MNase sequence in the pk19pAMNase vector (pk19pAMNase was a gift from Ulrich Laemmli, Addgene #86973) was replaced by the Apex2-6xHis sequence. Apex2 was amplified from the plasmid Apex-2NLS_pMT-Hygro ^28^ and inserted in the pK19 vector between EcoRI and BamHI restriction enzymes using the primers “pProA-Apex2-3Gly-6His FW” and “pProA-Apex2-3Gly-6His RE”. Cloning was performed with In-Fusion cloning kit (Clontech). The list of primers is available in the Supplementary Table S1.

### Expression and purification of pA-Apex2 enzyme

The bacterial expression plasmid (pk19pA-Apex2-6xHis) was transformed into Bl21DE3 Gold (Stratagene# 230132) according to manufacturer instruction. Transformed colonies were grown overnight and re-cultured in a fresh 1 liter of LB medium until the OD reaches to a value 0.6. The protein expression was then induced by adding IPTG (0.25mM) and cultured for additional 3 h. Cells were harvested and resuspended in 40ml of lysis buffer (25mM Tris pH: 7.2, 10% Glycerol, 150mM NaCl, 0.1% IGPAL, 1mM DTT, 10mM Imidazole, and protease inhibitors). Cells were at first lysed with lysozyme and then sonicated. The cell lysate was centrifuged at 45000 rpm for 30min at 4 °C. The cleared lysate was then incubated with 1ml of TALON metal affinity (Takara #Z5504N) resin according to manufacturer instruction. Lysate with the beads were passed through a column (Econo-Pack Disposable Chromatography Columns, 10 ml, BioRad) by gravity flow. The column was then washed three times with lysis buffer, and twice with wash buffer (25 mM Tris pH: 7.2, 10% Glycerol, 150 mM Nacl, 0.1% IGPAL, 1 mM DTT, 20 mM Imidazole, and protease inhibitors). The enzyme ProtA-Apex2 was eluted with 5 ml of wash buffer containing 200 mM Imidazol, collected in 1 ml fractions. A small aliquot of the collected fractions was loaded on to an SDS–PAGE gel and stained with InstantBlue Coomassie protein stain (abcam #ab119211) according to the manufacturer’s instruction. Fractions containing pA-Apex2 were pulled together and dialyzed overnight in dialysis buffer ((25 mM Tris pH: 7.2, 10% Glycerol, 150 mM NaCl, 0.1% IGPAL, 1 mM DTT, and protease inhibitors). The dialyzed protein solution was then concentrated using Amicon Ultra-4 Centrifugal Filter Units 30 K (Millipore, MA #UFC803024) according to manufacturer’s protocol. The enzyme was snap-frozen and stored at −80 °C in storage buffer (25 mM Tris pH: 8, 30% Glycerol, 150 mM NaCl, 0.1% IGPAL, 1 mM DTT, 2mg/ml of Biotin free BSA (Roth #0163.2), and protease inhibitors. The concentration of pA-Apex2 was determined using BSA standards.

### In-vitro activity assay of pA-Apex2

Cells were harvested, washed 2× in PBS and resuspended in hypotonic buffer (20 mM HEPES pH 7.9, 20 mM NaCl, 5 mM MgCl2, 1 mM PMSF, 1 mM DTT, cOmplete™ EDTA-free Protease Inhibitor Cocktail (Roche)) and incubated on ice for 10 min. Subsequently, cells were dounced with a 26^1^/_2_ G needle and again incubated on ice for 10 min. Nuclei were pelleted at 500 g, 5 min, 4°C and lysed in hypotonic buffer supplemented with 0.5% IGEPAL® CA-630 (Sigma). To the lysate, Benzonase (Millipore) was added and rotated at 4°C for 1 h. Subsequently, NaCl concentration was raised to 300 mM through addition of 5 M NaCl to lysate and rotated at 4°C for another 30 min. NaCl concentration was lowered back to 150 mM through addition of hypotonic buffer supplemented with 0.5% IGEPAL^®^ CA630 and lysate cleared by centrifugation (15000 g, 15 min, 4°C). For experiments shown in the supplementary Figures 1b, c, an aliquot of the sample was incubated with 5 μM pA-Apex2 and 500 μM Biotin-Phenol (BP) for 1 min. The biotinylation reaction was triggered by adding H_2_O_2_ to a final concentration of 2 mM for 1 min. The reaction was stopped by adding quenching solution. As a negative control similar reaction was performed either in absence of H_2_O_2_, BP, or H_2_O_2_ and BP.

### Western blotting

Biotinylated extract from In-vitro assay or biotin immunoprecipitated streptavidin bead bound proteins were eluted in Lemmli Buffer at 95°C and resolved on a SDS-PAGE gel (Serva #43264.01). Proteins were then transferred to a PVDF membrane using a Trans-Blot Turbo Transfer System (Bio-Rad) according to the manufacturer’s instructions. Membranes were blocked in blocking solution (2% Biotin free BSA in PBS) in a shaker for 1 hour at room temperature. Membranes were then incubated with HRP-Streptavidin (BioLegend #405210) (1:2000) in 2% Biotin free BSA in PBST (PBS with 0.1% Tween) for 1 hour in room temperature. The membranes were developed using Clarity™ Western ECL Substrate (BioRad #170-5061) and imaged using ChemiDoc™ Touch Imaging Syatem (BioRad).

### AMPL-MS proximity biotinylation and immunoprecipitation

For AMPL-MS experiments cells were grown in T75 flasks (Greiner) to a density of 1×10^7^ cells/ml. For each experiment 2×10^7^ cells were used, therefore 32×10^7^ for a set of experiment (i.e., including experiment, antibody control, biotin-phenol control, and H_2_0_2_ control). The cells were washed with 20 ml cold PBS in a 50 ml tube and centrifuged (Thermo scientific, Heraeus, Multifuge X3R) at 250 g, 4 °C for 5 min. Next, for the nuclear isolation the cell pellet was resuspended in three packed cell volumes (PCV) of Nuclear Isolation Buffer (20 mM Tris (pH7.6), 10 mM KCl, 2.5 mM MgCl_2_, 0.5mM EDTA, cOmplete™ Protease inhibitors), incubate on ice for 10 min. Following the incubation, the cell suspension was supplemented with NP40 to a final concentration of 1% and pass through a 20G needed. The nuclei were spun down at 500 g for 5 min, and washed with NIB with 0.1% NP40. The nuclei were then fixed using in AMPL-MS assay buffer (20 mM HEPES (pH 7.5), 150 mM NaCl, 2.5 mM MgCl_2_, 0.5mM EDTA, cOmplete™ Protease inhibitor, MG132, 0.5 mM Spermidine) containing 3.7% v/v PFA and incubated on a rotating wheel for 10 min in room temperature. The reaction was quenched by adding 1/20 volume of 2.5 M Glycine. Subsequently, the nuclei were washed twice with AMPL-MS assay buffer and briefly treated with H_2_O_2_ (5 mM) and quickly washed with assay buffer. Following the treatment, the nuclei were permeabilized with 0.25% TritonX100 for 6 minutes on ice and washed in assay buffer supplemented with 1% BSA (biotin free BSA). The nuclei were then blocked with Image-iT™ FX Signal Enhancer (Invitrogen™ #I36933) for 45 minutes on a rotating wheel at room temperature. The nuclei were then quickly washed with assay buffer and resuspended in antibody incubation buffer (assay buffer with 5% NGS (Jackson ImmunoResearch), 0.02% digitonin) and split equally into four 0.5ml low protein binding tubes. The respective antibodies were added (2.5 μg/reaction) and incubated overnight in the cold room on a rotating wheel. To remove unbound antibody, following day, the nuclei were washed thrice with wash buffer (assay buffer with 0.1% Tween 20). Next, the nuclei were incubated with 2.5 μg of pA-Apex2 in antibody incubation buffer for 2 hours in the cold room and another one hour in room temperature on a rotating wheel. After the incubation the nuclei were washed thrice with wash buffer to remove the unbound enzyme. Following the washes biotinylation was performed. The nuclei were first incubated with biotin tyramide (500 μM) in assay buffer for 20 min in room temperature in the dark, then H_2_O_2_ (1 mM) was added for 4 min to start the biotinylation reaction. The quenching buffer (Trolox-5 mM, Sodium Ascorbate-10 mM, Sodium Azide 10 mM) was added to stop the reaction. The nuclei were washed twice with wash buffer with quenching solution. A small amount of the nuclei was saved for immunofluorescence. The rest of the nuclei were subjected to lysis and decrosslinking by heating at 99°C on a Thermo shaker (800 rpm) for 1 hour in nuclear lysis buffer (100 μl PBST (PBS with 0.1% Tween 20) 30 μl of 10% SDS and 20 μl of 10% Sodium deoxycholate). To adjust the concentration of the detergent for immune precipitation the volume of the samples was made up to 1 ml with PBST. Subsequently the samples were treater with 100 U of Benzonase^®^ (Milipore #1.01654.0001) and spun down at 14000 rpm for 20 min at 4°C. A small fraction (5%) of the supernatant was kept aside as input. The rest of the lysate was incubated with 50 μl of precleaned streptavidin beads (Invitrogen, Dynabeads™ M-280 Streptavidin #11206D) at room temperature for 2 hr.

The beads were then washed twice with PBST, twice with PBST+1 M NaCl, twice with PBS. The beads were transferred to a new tube and washed thrice with 50 mM ammonium bicarbonate. A small fraction (5%) of the beads were saved for immunoblot analysis. The rest of the beads were subjected to on-bead digestion for mass spectrometry.

### Immunofluorescent staining

Immunofluorescent experiments were performed as described previously ^28^ with minor modifications. Briefly, the nuclei from AMPL-MS assays were adhered on a Poly-L-Lysine coated glass coverslip, washed with PBST with quenching solution. After washes the nuclei were incubated with secondary antibodies for respective bait coupled to Alexa Fluor 488, and anti-biotin antibody [Streptavidin, Alexa Fluor™ 555 #S32355, (1:600)] for 1h at room temperature (RT). Slides were again washed 3× 5 min with 0.1% Triton X-100/PBS and incubated with DAPI for 3 min. Excess DAPI was washed off with 0.1% Triton X-100/PBS for 5 min and samples mounted with VECTASHIELD (Vector Labs). Images were acquired using Leica TCS SP8 Confocal Microscope, processed and quantified using Fiji^62^.

### Mass spectrometry

Mass spectrometry based proteomic experiments were performed as described previously (*Kochanova et. all*.) with minor modifications. Briefly, beads were washed three times with 50 mM NH_4_HCO_3_ and incubated with 10 ng/μL trypsin in 1 M urea 50 mM NH_4_HCO_3_ for 30 minutes, washed with 50mM NH_4_HCO_3_ and the supernatant digested overnight in presence of 1mM DTT. Digested peptides were alkylated and desalted prior to LC-MS analysis. The peptide mixtures were subjected to nanoRP-LC-MS/MS analysis on an Ultimate 3000 nano chromatography system coupled to a Qexactive HF or a Orbitrap Exploris-480 mass spectrometer (both Thermo Fisher Scientific) in 2–4 technical replicates (5 μl each).

### Database search

MaxQuant 1.6.1.4^63^ was used to identify proteins and quantify by LFQ with the following parameters: Database, dmel-all-translation-r6.08.fasta (Flybase); MS tolerance, 10 ppm; MS/MS tol, 20 ppm; Peptide FDR, 0.1; Protein FDR, 0.01; Min. peptide Length, 5; Variable modifications, Oxidation (M); Fixed modifications, Carbamidomethyl (C); Peptides for protein quantitation, razor and unique; Min. peptides, 1; Min. ratio count, 2. Match-between-runs (MBR) option was selected. Technical replicates were assigned to one experiment (biological replicate). Experimental and control samples (treated with biotin-phenol and DMSO, respectively) were loaded into the same MaxQuant run. Samples from different cell lines and time points were run separately.

### Data analysis

The output files from MaxQuant (proteinGroups.txt) were analyzed in R environment. Data was filtered such that proteins that are present in 2 of the 3 replicates of at least one condition were taken for the analysis. Following filtering, MinProb imputation algorithm with q = 0.01 was performed to impute the missing values and limma (CIT) based differential expression analysis was carried out. Proteins were considered significant if the LFC > 1 and FDR ≦ 0.05. Over Representation Analysis was performed using the enrichGO function from clusterProfiler (CIT) ^49^package v3.12.0 by taking the significant proteins (FDR ≦ 0.05 cut-off for predicted GO terms) and without a background set. Corresponding GO plots were also generated with R environment. For the tree plot, GO terms were clustered based on GO semantic similarity. Based on the representation in each cluster, a summarized GO term was the written. Scripts used for the analysis can be provided on request.

### Plots and statistical analysis

All statistical analysis was performed in R environment, except for Fig. 1c and 2b. Plots and graphs were generated in R environment, except for Fig..1c, 2b, 2g, 3f, 3e, 3g and 3h. All schematic figures were created with BioRender.com.

### Data sources

The datasets produced in this study are available in the ProteomeXchange Consortium via the PRIDE ^64^ partner repository with the identifiers: PXD044295 (Proteomics) and PXD044296 (Histone PTMs).

### Histone PTM analysis

To look at histone modification associated with different chromatin domains as shown in Figure: 2f, and supplementary Figure: 3g, h, i., we performed AMPL-MS assay as explained above and performed histone PMT analysis as described previously ^65^ with minor modifications. In brief, the biotinylated proteins were immunoprecipitated using streptavidin beads as earlier and eluted in 1xSDS lysis buffer. The proteins were then resolved on a precast SERVAGel™ TG PRiME™ 4-20 % (SERVA Electrophoresis GmbH) gel and stained with InstantBlue Coomassie Protein Stain. Protein bands which correspond to the histones (expected between 11-17 kDa) were cut out and destained. Following destaining, in-gel histone acylation using propionic anhydride and digestion with trypsin [MS Grade Pierce™ Trypsin Protease (Thermo Scientific)] were performed. The histone peptides were then extracted and cleaned using a C8 Stagetip ^66^ before mass spectrometry analysis.

## Acknowledgements

We would like to thank the members of the Imhof lab and Becker department for their lively discussion and excellent suggestions. In addition, we thank Markus Hohle from QBM for his constant support. Work in the AI lab was funded by grants from the DFG and QBM, grant numbers 213249687 (CRC1064) 325871075 (CRC1309) and 419067076 (SPP2191).

## Author contributions

RC and AI conceptualized the project. RC performed the experiments with validation from JH. IF performed LC/MS measurements. MB performed the histone modification analysis. AVV performed data analysis for all mass spectrometry data and prepared the corresponding figures with suggestions from RC. RC and AI wrote the manuscript with comments from all authors. All schematic figures were created with BioRender.com.

